## Supplementary information for "The increase in cell volume and nuclear number of the koji-fungus *Aspergillus oryzae* contributes to its high enzyme productivity"

Supplementary materials
Figure S1-5.
Table S1-7 legends.
Movie 1-6 legends.

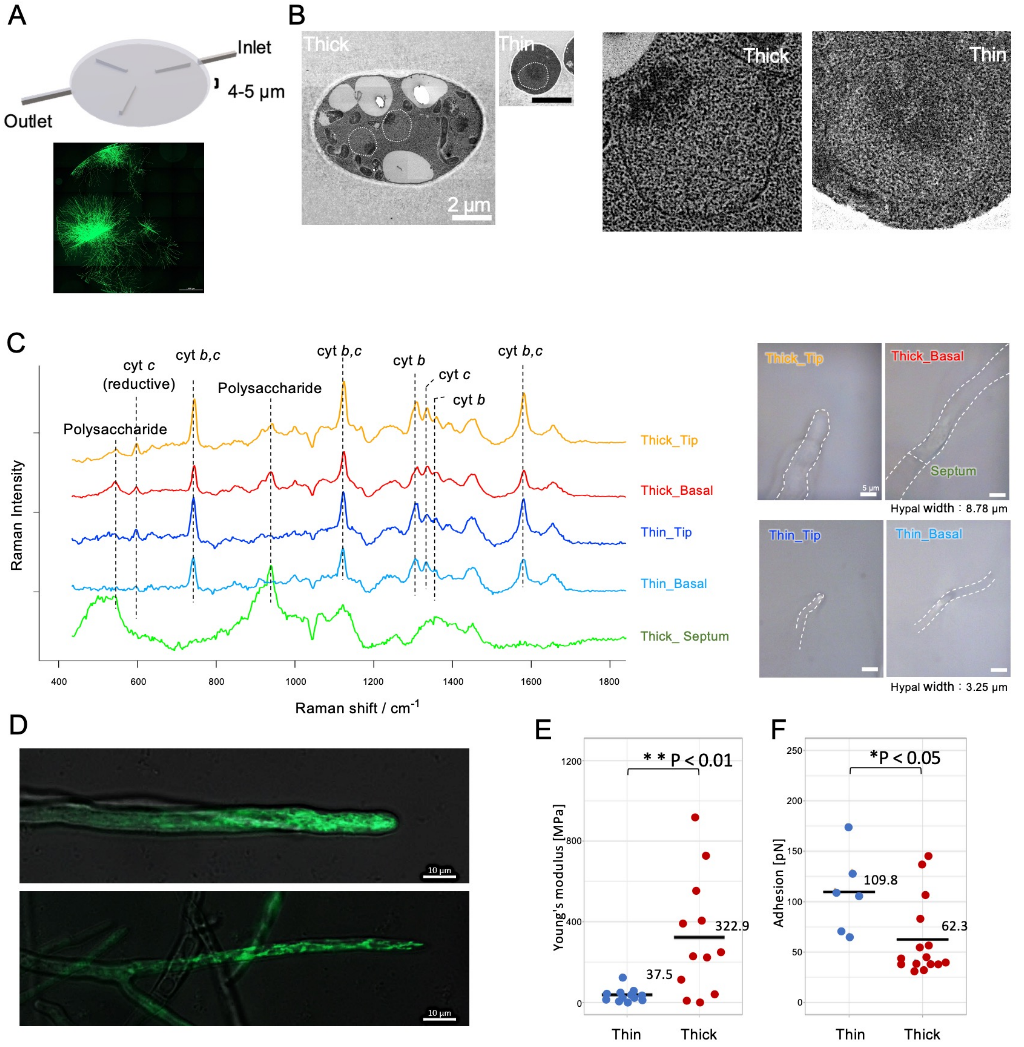


Fig. S1. (A) Microfluidic device for 2D observation of hyphae (upper) and overall image of *A. oryzae* RIB40 expressing H2B-GFP cultured within the device (lower). (B) Cross-sectional TEM images. White dotted lines indicate nuclei (left). Enlarged view of the nuclear region (right). Scale bar: 2 μm. (C) Averaged Raman spectra (left) measured at different areas of thick and thin hyphae (right). (D) Mitochondrial staining using Rhodamine 123 in thick hyphae (upper) and thin hyphae (lower). (E) Young's modulus of hyphal tips measured by AFM. (F) Adhesion properties of hyphal tips measured by AFM.

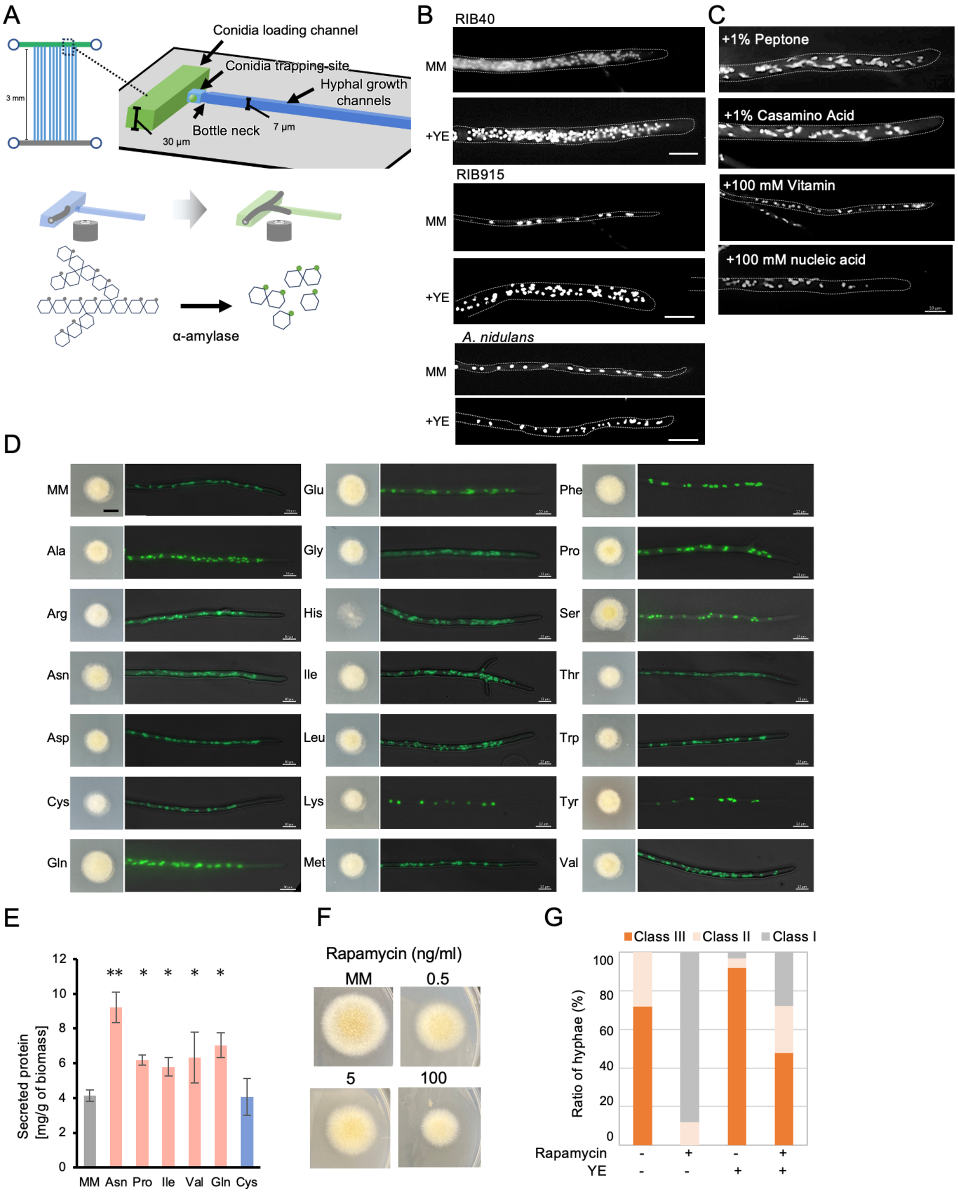


Fig. S2. (A) Microfluidic device for isolating single hyphae and measuring enzymatic activity. The design was modified from a previous study (31). α-amylase hydrolyzes the starch backbone, releasing a fluorescent signal. (B) Hyphal images stained with SYBR Green of *A. oryzae* RIB40, RIB915, and *A. nidulans* grown for 3 days on the minimal medium with or without 1% yeast extract. Scale bar: 20 µm. (C) Hyphal images stained with SYBR Green of *A. oryzae* RIB40 grown for 3 days on the minimal medium with 1% peptone, 1% casamino acid, 100 µM vitamin Bs or 10 mM nucleic acid. Scale bar: 10 µm. (D) Colonies and hyphae stained with SYBR Green of *A. oryzae* RIB915 grown for 3 days on the minimal medium with 0.1% amino acid respectively. Scale bars: colony, 1 cm; hyphae, 10 µm. (E) Secreted protein per biomass in RIB915 cultured in the minimal medium with 0.1% amino acid (mean ± S.E., n=3, ** $p < 0.01$, * $p < 0.05$, t-test). (F) Colony morphology of RIB40 cultured for 3 days on the minimal medium containing 0, 0.5, 5, or 100 ng/ml rapamycin. (G) Ratio of class I-III hyphae in *A. oryzae* RIB40 cultured on minimal medium with or without 100 ng/ml rapamycin and 1% yeast extract (n=50).

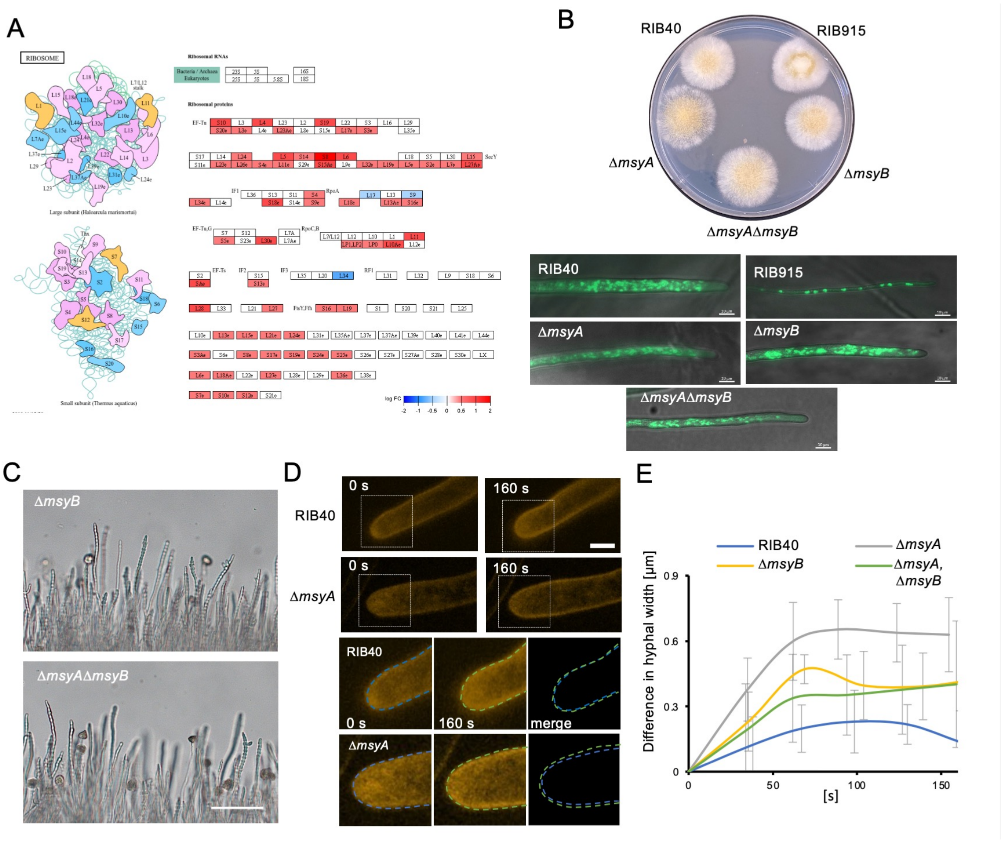


Fig. S3. (A) KEGG PATHWAY visualization showing changes in ribosome-related gene expression in thick hyphae compared to thin hyphae. (B) Colonies and hyphae stained with SYBR Green of *A. oryzae* RIB915, RIB40, Δ*msyA*, Δ*msyB*, and Δ*msy*AΔ*msyB* cultured in minimal medium for 3 days. Scale bar: 10 μm. (C) Images of hyphae of Δ*msyB* and Δ*msy*AΔ*msyB* strains after 3 min of low osmotic stress. Scale bar: 200 μm. (D) Images of hyphal tips stained with FM4-64 of RIB40 and Δ*msyA* immediately after low osmotic stress (0 s) and 160 s later (upper). Scale bar: 5 μm. Enlarged images of the same hyphal tips (lower), showing outlines at 0 s in blue and at 160 s in green. (E) Time course of differences in hyphal width every 30 s for 150 sec (mean ± S.E., n=3).

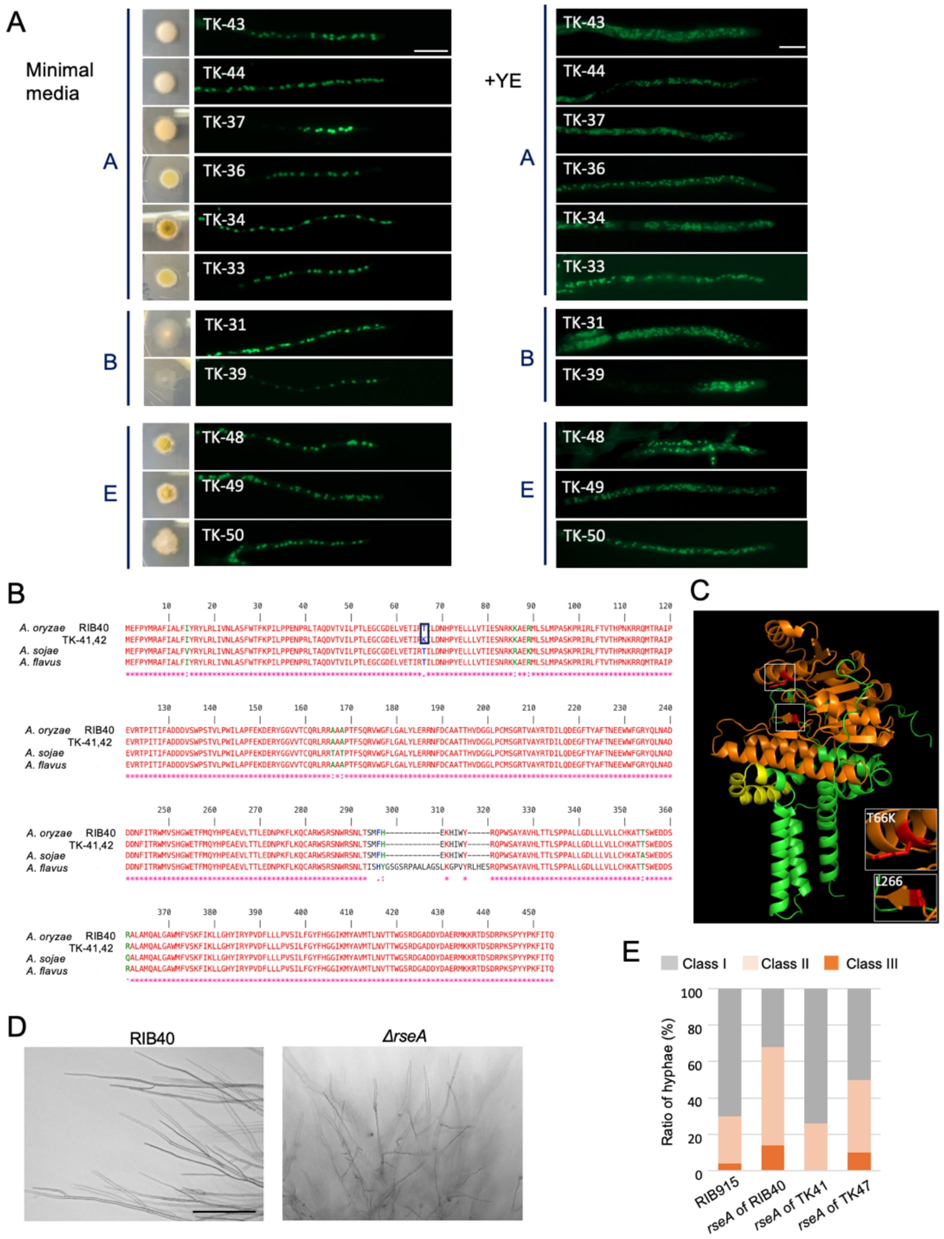


Fig. S4. (A) Colonies and hyphae stained with SYBR Green of strains from clades A, B, and E cultured on the minimal medium (left) with yeast extract (right) for 3 days. Scale bar: 20 μm. (B) Alignment of *rseA* orthologs in *A. oryzae* RIB40, TK-41, TK-42, *A. sojae*, and *A. flavus*. Amino acid substitution T66K in *A. oryzae* is highlighted in black. (C) Predicted structure of *A. oryzae* RseA by AlphaFold2. Orange indicates the Glycosyltransferase-like family 2 domain, and yellow indicates a membrane-bound protein region predicted by InterPro. T66K and L266 are highlighted in red. (D) Hyphae at the colony periphery of RIB40 and Δ*rseA* cultured on the minimal medium for 3 days. Scale bar: 300 μm. (E) Ratio of class I-III hyphae in RIB915 and the strain expressing *rseA* of RIB40 (with increased nuclei), TK-47 (with increased nuclei), or TK-41 (without increased nuclei) cultured on the minimal medium for 3 days.

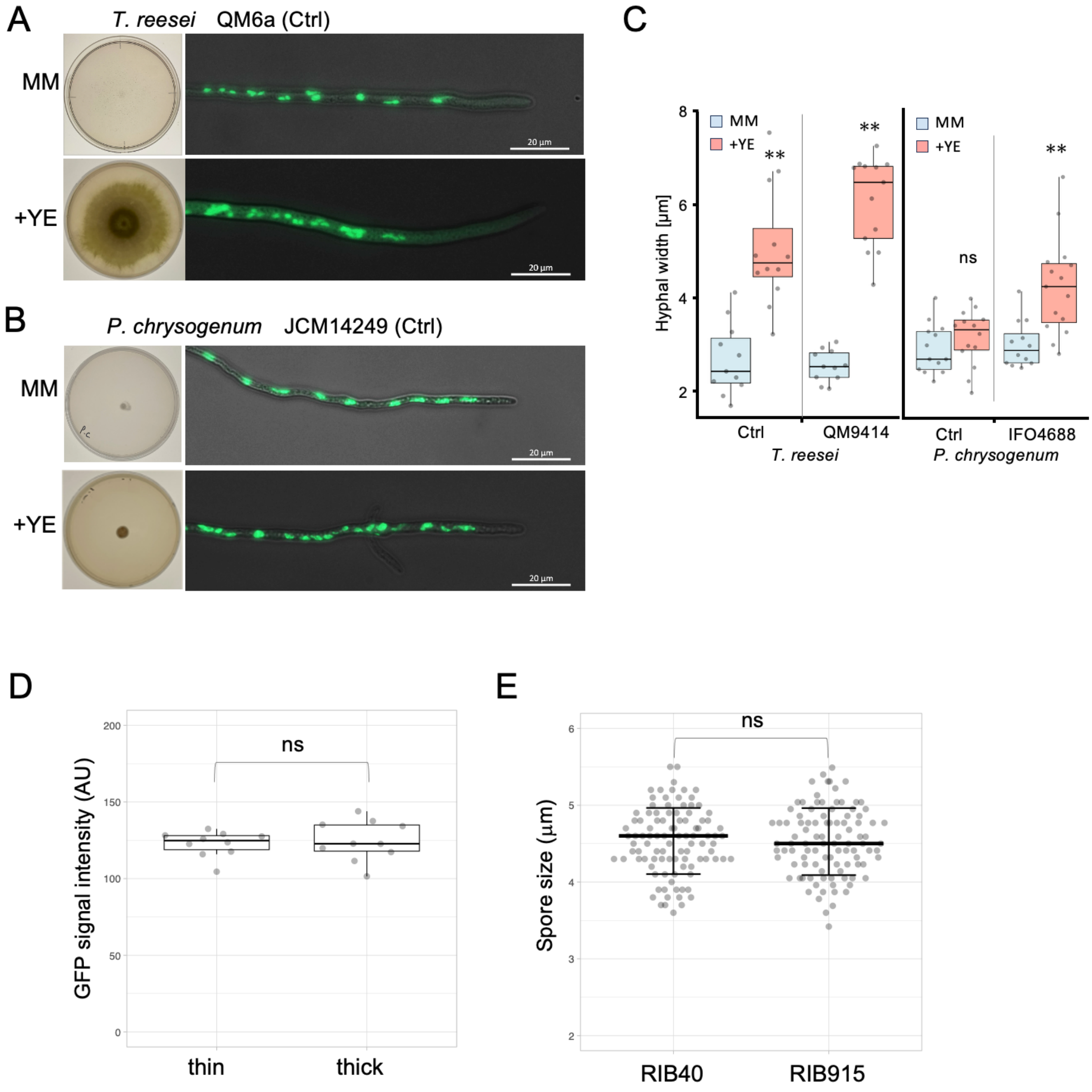


Fig. S5. (A, B) Colonies and hyphae stained with SYBR Green of the *T. reesei* control strain (QM6a) (A) and the *P. chrysogenum* control strain (JCM14249) cultured on the minimal medium with or without 1% yeast extract for 3 days. Scale bar: 20 µm. (C) Box plots of hyphal width of industrial strains and their controls under the same conditions in G (n=11–15, ** $p < 0.01$, * $p < 0.05$, t-test). (D) Box plots of signal intensity of GFP-histon in *A. oryzae* RIB40 (n=10). (E) Box plots of spore size in *A. oryzae* RIB40 and RIB915 (n=100).

Table S1. Annotated data of RNAseq in *A. oryzae* RIB40 and RIB915, and *A. nidulans* grown in the minimal medium with or without yeast extract.

Table S2. Annotated data of RNAseq in *A. oryzae* RIB40 thick or thin hyphae.

Table S3. Annotated data of RNAseq common in upregulated gene in *A. oryzae* RIB915 grown with yeast extract and in *A. oryzae* RIB40 thick hyphae.

Table S4. SNP analysis of ORFs in Clades F between TK-32 and TK-38.

Table S5. SNP analysis of ORFs in Clades G between TK-41 and TK-47.

Table S6. Strains used in this study.

Table S7. Composition of minimal medium.

Movie 1. 3D images of hyphae without increased nuclei (left) and with increased nuclei (right) in *A. oryzae* RIB40. Each nucleus is indicated with different colors by Imaris soft. Scale bar: 5 µm.

Movie 2. Emergence of thick hyphae with increased nuclei by branching in *A. oryzae* RIB40 expressing H2B-GFP. Scale bar: 20 µm. Elapsed time is indicated in min.

Movie 3. Successive nuclear division within the newly emerged thick branched hypha in *A. oryzae* RIB40 expressing H2B-GFP. Scale bar: 10 µm. Elapsed time is indicated in min.

Movie 4. Mycelial growth of hyphae with increased nuclei (green) and without increased nuclei (white). Scale bars: 200 µm. Elapsed time is indicated in h.

Movie 5. Amylase activity was monitored with a fluorescent substrate under conditions where a single thick or thin hypha grew in a microfluidic channel. Scale bar: 10 µm. Elapsed time is indicated in min.

Movie 6. Each hypha was dissected using laser microdissection and collected separately.
