## Supplementary material for "The increase in cell volume and nuclear number of the koji-fungus *Aspergillus oryzae* contributes to its high enzyme productivity": Table S7

Table S7. Composition of Minimal medium

| Minimal Medium | | |
|---|---|---|
| Glucose | 10 | g |
| $NaNO_3$ | 6 | g |
| $KH_2PO_4$ | 1.52 | g |
| KCl | 0.52 | g |
| $MgSO_4 \cdot 7H_2O$ | 0.52 | g |
| Hunter's Trace element | 2 | mL |
| pH | 6.5 | |
| per litter | | |

| Hunter's Trace element | | |
|---|---|---|
| $ZnSO_4 \cdot 7H_2O$ | 2.2 | g |
| $H_3BO_3$ | 1.1 | g |
| $MnCl_2 \cdot 4H_2O$ | 0.5 | g |
| $FeSO_4 \cdot 7H_2O$ | 0.5 | g |
| $CoCl_2 \cdot 6H_2O$ | 0.16 | g |
| $CuSO_4 \cdot 5H_2O$ | 0.16 | g |
| $(NH_4)6Mo7O_{24} \cdot 4H_2O$ | 0.11 | g |
| per 100 mL | | |
